# Kinetic characterization of the interaction of NO with the S_2_ and S_3_-states of the oxygen-evolving complex of Photosystem II

**DOI:** 10.1101/2021.01.10.426130

**Authors:** Gert Schansker

## Abstract

The reactivity of the S_3_ and S_2_ states towards NO and NH_2_OH was studied and compared using the period-4 oscillations in the F_0_-value induced by a train of single turnover Xenon flashes spaced 100 ms apart to monitor the reaction kinetics. The flash frequency also determined the time resolution of the assay, i.e. 100 ms. The S_2_ and S_3_-states were created by one and two single turnover pre-flashes, respectively. The NO-concentration-dependence of the S_3_-decay indicated that at low NO-concentrations an S_2_-state was formed as an intermediate, whereas at higher concentrations a seemingly monophasic decay to the S_1_-state was observed. The sigmoidal concentration dependence indicated that a fast interaction of the S_3_-state with (at least) two NO-molecules is necessary for the fast S_3_ to S_1_ decay (τ ~0.4 s at 1.2 mM NO). The pH-dependence of the S_3_-decay suggests that a protonation-reaction (pK ~6.9) is involved in the S_3_ to S_1_ decay. At intermediate NO-concentrations the protonation is only partially rate limiting, since the pH effect is more pronounced at high compared to intermediate NO-concentrations. A comparison of the reactivity of NO and hydroxylamine suggests that hydroxylamine reacts more efficiently with the S_1_ and S_2_ states, whereas NO reacts more efficiently with the S_3_-state. Based on our present knowledge of the oxygen evolving complex a possible reaction mechanism is proposed for the interaction between NO and the S_3_ state.

## Introduction

The ability of photosystem II (PS II) of eukaryotic photosynthetic organisms to break two water molecules into an oxygen molecule and four protons while extracting four electrons, at room temperature, has drawn much attention and many extremely sophisticated techniques have been applied to elucidate the chemical basis for this reaction. For recent reviews see Yano and Yachandra 2014, Vinyard and Brudvig 2017, Junge 2019, Lubitz et al. 2019. Here, I will limit myself to the literature that is of direct relevance to the presented experiments. These experiments were carried out in the period 1999-2000. That is a long time ago, but since no other studies on kinetics of the interaction between NO and the S_2_ and S_3_ states of the oxygen evolving complex at room temperature have been published since then, publication still has value. The reactivity of NO towards the S_2_ and S_3_ states was studied in detail making use of the period-4 oscillations in the F_0_-signal induced by a train of single turnover flashes spaced 100 ms apart that were preceded by one or two pre-flashes to induce either the S_2_ or S_3_ state of the oxygen evolving complex. By varying the time between pre-flashes and flash train it is possible to follow the decay kinetics of the created S-state. These data were compared with those of NH_2_OH (hydroxylamine).

Around 1990, Johannes Messinger published a series of articles studying the reactivity of NH_2_OH and NH_2_NH_2_ (hydrazine) towards the S_2_ and S_3_ states using a bare oxygen electrode and analyzing the period-4 oscillations in the flash-induced O_2_ yield (a.o. Messinger and Renger 1990, 1994, Messinger et al. 1991). The fluorescence method has several major advantages over the bare oxygen electrode. The time resolution is considerably higher. In this study a maximum flash frequency of 10 Hz was used with which a time resolution of 100 ms was obtained, but it is technically possible to increase the flash frequency further (cf. Ananyev and Dismukes 2005) and thereby the time resolution. A second advantage is that a cuvette-based configuration was used and the cuvettes were not sensitive to corrosive gases like NO. It is also much easier to treat a sample in a cuvette with a gas by bubbling than a sample on a bare oxygen electrode. There are also a few disadvantages. The fluorescence signal is a complex signal, which gives rise to a high level of versatility but at the same time makes interpretation more complicated. The interpretation of the oxygen yield is in comparison with the fluorescence signal extremely straightforward. Another disadvantage in the second half of the nineties was that it was possible to buy bare oxygen electrodes, but there was no commercial equipment for the fluorescence signal. I worked, therefore, with a homebuilt apparatus based on a design of David Kramer. At the moment this is less of a problem. The MULTI-COLOR-PAM with which I work at my present employer (Heinz Walz GmbH) is capable of the detection of period-4 oscillations and also Photon Systems Instruments (PSI) sells an instrument that according to the specifications is capable of measuring period-4 oscillations.

In mechanistic terms the period-4 oscillations in the F_0_ signal are somewhat special. The period-4 oscillations in the F_M_-signal can be explained on the basis of the S-state dependent lifetime of P680^+^, a strong fluorescence quencher (Zankel 1973, Steffen et al. 2005). Hundred ms after a single turnover flash the concentration P680^+^ is essentially zero and cannot explain the S-state dependent oscillations in F_0_ that have a phase opposite to those observed for the oscillations in the F_M_ level (cf. Delosme and Joliot 2002, Fig. 5). The mechanistic cause of these oscillations has not been elucidated yet. An intriguing observation in this respect was made by Delrieu (1993), who noticed that the oscillations in the F_0_-signal were sensitive to the antenna size. The advantage of the period-4 oscillations in the F_M_ level is that they are larger, but these oscillations also suffer more from the period-2 oscillations related to the acceptor side of PS II (Robinson and Crofts 1987, Shinkarev and Wraight 1993). However, due to the equilibrium between Q_A_ and Q_B_ (Diner 1977) the period-4 oscillations in the F_0_-signal are also not entirely free from contributions of the period-2 oscillations.

The data in this study were collected in the laboratory of Vasili Petrouleas (Demokritos, Greece) where I worked at the time as a Post-doc. The interest in the lab changed in this period from the acceptor side to the donor side of PS II. This change was strongly stimulated when a PhD student in the lab, Harris Goussias, discovered how you could make a new EPR-multiline by incubating PS II-membranes at −30 °C with NO (Goussias et al. 1997, Ioannidis et al. 1998). This multiline later was found to represent the super-reduced S_-2_ state (Schansker et al. 2002). Flash-oxygen and flash-fluorescence measurements are ideal for such determinations, but it became quite quickly clear that the quality of the samples was very important as well. The high quality of the NO-treated samples that Josephine Sarrou made for me were in this respect critical. The much better quality of the dataset in this study compared the data presented in the Proceedings of the 1998 Budapest meeting (Schansker and Petrouleas 1998) is to a large extent due to improvements in the isolation of the PS II membranes around 1998 from which not only the EPR-work profited.

The other half of the story is the treatment of those period-4 oscillations in the F_0_ signal. As shown by Kurreck et al. (2000), PS II membranes suffer from severe PQ-pool quenching. FeCy was used to keep the PQ-pool in the oxidized state, however, at a flash frequency of 10 Hz, FeCy could not keep up with the rate at which the PQ-pool became reduced and an F_0_ fluorescence rise was still observed. It is possible, and no doubt elegant, to create a baseline fit and to subtract the baseline. Here, a different approach was chosen. Delrieu (1998) had forced the minima to zero and we did the same thing for the maxima, which were forced to 1. In addition, a second treatment was applied to the data. From the beginning the signal had been sensitive to a 50 Hz oscillation related to the mains. This problem seemed unsolvable. Petrouleas proposed to eliminate this problem by using a gliding two-point average. This eliminated at the same time any potential period-2 oscillation caused by the PS II acceptor side. Later on, Laszlo Sasz, a collaborator of Imre Vass at the Biological Research Center in Szeged (Hungary), came with a different solution for the mains problem. He wrote new software for the instrument in which the F_0_-values accumulated over 20 ms (one 50 Hz cycle) were averaged. But even then, there remained some residual PS II acceptor side related period-2 oscillations, which, as noted above, were due to the equilibrium between Q_A_ and Q_B_. As a consequence, the gliding two-point average was used throughout this study. It may be argued that these data manipulations distort the data, but there it can be noted that with respect to the S_3_ to S_1_ decay at higher NO-concentrations, it could always described perfectly by mono-exponential kinetics.

Another technical problem had to do with the NO gas. Nitrogen gas was led through the cuvette in order to bring down the oxygen concentration and since the length of the experiments was quite short, we assumed initially that the NO concentration would be stable during the time of the experiment. However, this turned out to be an incorrect assumption. This led to the creation of two datasets. One series where the experiment was started immediately after the NO treatment and where the equilibration time between NO and sample was very short and a series where the sample was incubated for 1 minute with NO before the experiment was started.

The study has a methodological aspect in the sense that it discusses and illustrates different aspects of the period-4 oscillations measured on PS II membranes. In this way, I hope that period-4 oscillations in the F_0_ signal can be an interesting tool for the study of the reactivity of the S-states towards exogenous electron donors as well as the decay kinetics of the S-states under a variety of conditions.

With respect to the kinetic data themselves, an attempt is made to explain the interaction of NO with the S_3_ state on the basis of our present understanding of the catalytic cycle of the oxygen evolving complex. NO is known to interact more efficiently with radicals (e.g. Eiserich et al. 1995, Ford 2004) than with metals (e.g. Ford 2004), but the present consensus is that on each S-state transition, including the S_2_ to S_3_ transition, manganese is oxidized (Dau et al. 2012, Siegbahn 2012, Yano and Yachandra 2014, Vinyard and Brudvig 2017, Junge 2019, Lubitz et al. 2019). At the same time it is now thought that on the S_2_ to S_3_ transition another water molecule binds to the manganese cluster. It is proposed that the kinetic data can be explained assuming that the combination of an electron donation by NO and a protonation can reverse this water binding and that the intermediate state produced this way, which is not resolved kinetically, can react with a second NO molecule after which the OEC ends up in the S_1_ state. At low NO concentrations this hypothetical intermediate state decays to the S_2_ state.

## Material and Methods

### Photosynthetic material

PS II-membranes were prepared from spinach according to (Berthold et al. 1981) with some minor modifications. The chlorophyll concentration was determined according to Arnon (1949). For the measurements the samples were diluted to 10 μg Chl ml^−1^ in a buffer containing 0.4 M sucrose, 15 mM NaCl, 5 mM MgCl_2_ and 40 mM MES (at pH-values below 6.8), 40 mM HEPES (at pH-values above 7.2) or 20 mM of each buffer (at intermediate pH-values).

### Fluorescence meter

Fluorescence measurements were carried out using a home-built fluorescence apparatus based on a design of dr. DM Kramer (Kramer et al. 1990). Saturating single turnover flashes were produced by a Xenon-lamp (EG&G FX-1248) shielded by a 1 mm KG5 filter. The time interval between the flashes in the presented experiments was 100 ms. Measuring flashes spaced 50 μs apart were produced by an array of 8 LEDs (Hewlett Packard HLMA-CH00). The LEDs were turned on, initially, 1 ms before each flash and later 20 ms beforehand; they were turned off 5 ms after the flash. The fluorescence intensity was measured using a photodiode (Hamamatsu S2744-08). A cut off filter RG695 (1 mm) was placed in front of the photodiode to protect the diode against actinic light from the flash lamp. The collected data were converted using a 16 bits AD7885 analog to digital converter, which was synchronised with the measuring flashes. The interface card was home made.

### Measurement of the period-4-oscillations

For the measurements a miss rate of about 8% was observed (Ioannidis et al. 2000), which was probably due to recombination reactions between flashes (Lavergne and Rappaport 1998); double hits occurred in about 4% of the centers. Both processes caused damping of the oscillations in the F_0_’-level (see e.g. Fig. 1A). In the samples a small fraction of centers in which no forward electron transfer towards Q_B_ occurred and in which re-oxidation of Q_A_^−^ could only occur via recombination with the donor side was observed (Schansker et al. 2002). In these centers no re-oxidation of Q_A_ occurred between flashes causing an F_0_’-jump on the first flash. The different S-states modulate the F_0_’-level: in the S_0_ and S_1_ states the F_0_’ level is lower than in the S_2_ and S_3_ states (e.g. Joliot et al. 1971, Delosme and Joliot 2002, Schansker et al. 2002).

**Fig. 1.**
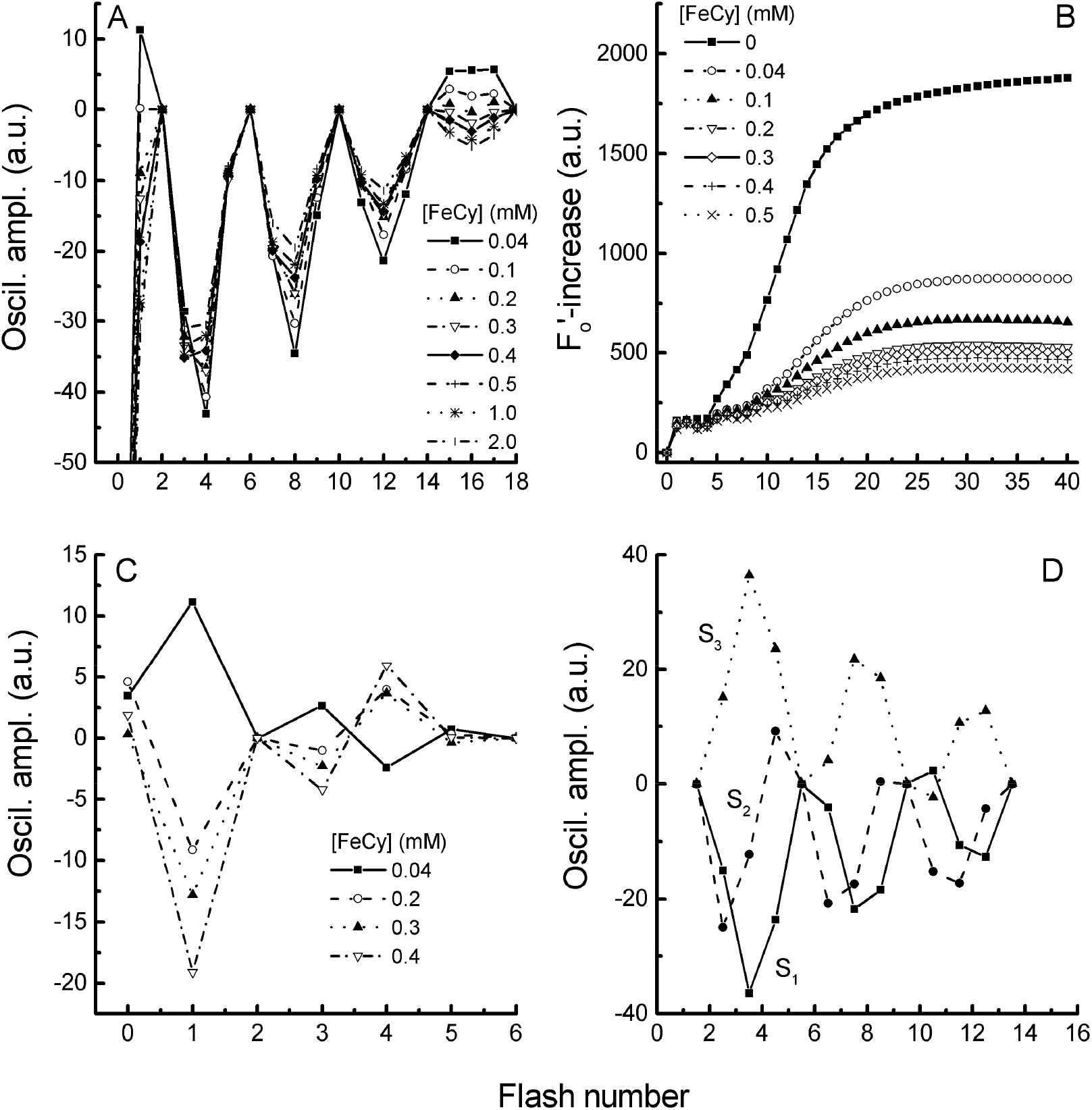
Effect of FeCy on the period-4-oscillations in the F_o_’-level. A. Relationship between [FeCy] and the depth of the period-4-oscillations; B. Ability of various FeCy-concentrations to keep the plastoquinone pool oxidized during a series of flashes spaced 100 ms apart; C. Inversion of period-2-oscillations by a one minute incubation of PS II-membranes with various FeCy-concentrations, the oscillations in the presence of 0.1 mM FeCy were subtracted from the oscillations measured at the indicated FeCy-concentrations; D. Form of the period-4-oscillations of the different S-states.

### Plastoquinone pool

PS II-membranes have a smaller plastoquinone pool than thylakoid membranes. Plastoquinone pool limitations increase the number of misses. To limit this, FeCy was added to keep the plastoquinone pool oxidized. FeCy quenches variable fluorescence to a considerable extent but has little effect on F_0_ (Murata et al. 1973, Butler and Kitajima 1975). Fig 1A demonstrates that only high FeCy-concentrations cause some quenching of the period-4-oscillations. FeCy was also used by Delrieu and Rosengard (e.g. Delrieu and Rosengard 1987, 1991) in their studies of the period-4-oscillations in the fluorescence signal.

If no electron acceptor is added the F_0_’-level increases strongly after a few flashes (Fig. 1B). This increase is unrelated to a built up of Q_A_^−^ (Schansker and Petrouleas 1998) but is related to plastoquinone pool quenching (e.g. Amesz and Fork 1967, Vernotte et al. 1979). In PS II-membranes static quenching by oxidized plastoquinone is 8-times stronger than in thylakoid membranes (Kurreck et al. 2000).

For the NO-experiments a time-resolution of at least 100 ms is necessary (Schansker and Petrouleas 1998, Ioannidis et al. 2000). At this flash frequency FeCy cannot keep the plastoquinone pool fully oxidized (Schansker and Petrouleas 1998 and Fig. 1B) and as a consequence a sigmoidal increase of the F_0_’-level is observed. For the analysis of the period-4-oscillations this increase was corrected for (see below).

### Period-2-oscillations

Period-2-oscillations are associated with electron transfer from Q_A_ to Q_B_ (e.g. Robinson and Crofts 1987, Shinkarev and Wraight 1993). This process has a halftime in the hundreds of microseconds. As is demonstrated in Fig. 1C period-2-oscillations can also be observed on much longer timescales. By incubating the samples with increasing concentrations of FeCy these period-2-oscillations can be inverted (Fig. 1C). Electron transfer between Q_A_ and Q_B_ is reversible (e.g. Diner 1977). The equilibrium constant for this reaction is pH-dependent (Crofts et al. 1984). At a more alkaline pH, the equilibrium is shifted more towards Q_A_. Therefore, oscillations in the Q_B_^−^ concentration with a period-2 will also induce period-2 oscillations in the Q_A_^−^ concentration that are stronger at more alkaline pH-values. Because the period-4-oscillations are small, small acceptor side dependent period-2-oscillations may affect the measurements. Fig. 1C also shows that the period-2-oscillations are dampened within 6 flashes and that incubation of samples for 5 minutes with 100 μM FeCy minimizes the period-2-oscillations.

Even so, the need was felt to find a way to eliminate the period-2-oscillations from the traces in a reliable way. An effective way to eliminate the period-2-oscillations was to add each pair of measuring points together and to divide the sum by 2 (gliding point average). In Ioannidis et al. (2000) this correction-method is demonstrated.

### Determination of the decay times (τ)

Rate constants for the interaction between NO and the S-states were determined in the following way: The time between the pre-flashes and the flash series was varied between 0.1 s and a few seconds. The measured period-4-oscillations were corrected for the baseline and period-2-oscillations as described in Ioannidis et al. (2000). For the S_3_-decay, the time dependence of the average oscillation depth of flashes 3 and 4 (about 15 time points were measured) was fit with 1 or 2 exponentials. The S_2_-decay-time was determined in the same way, but in that case the time-dependence of the average of flashes 4 and 5 was used for the analysis. In Fig. 5 the standard deviation of the halftimes determined at different pH-values is plotted. At higher pH-values the standard deviation is larger because the period-4-oscillations are smaller. In Fig. 1D it is shown what the period-4-oscillations of the individual S-states look like after correction.

### Chemical treatments

*NO* - In the presence of oxygen, NO reacts to NO_2_, which is able to destroy PS II-membranes. This reaction was minimised by using an air-tight cuvette and by bubbling the sample in the cuvette with 20 ml of N_2_-gas 5 minutes before the sample was bubbled with 20 ml of the NO/N_2_ mixture. NO may react with residual oxygen but the NO_2_ produced will be bubbled out during the procedure, limiting in this way the damage to the PS II-membranes. Bubbling of photosynthetic material with gas is thought to be bad for Photosystem II. In my hands the damage seemed to be limited: the period-4-oscillations that depend on an intact donor side could still be observed and the measurements were quite reproducible. The major effect of the bubbling with either N_2_ or NO seems to be that the plastoquinone pool becomes more reduced, causing an up-shift of the F_0_’-level. That this up-shift is related to the reduction state of the plastoquinone pool is indicated by the observation that the maximum F_0_’-level (all plastoquinone reduced) was not changed (not shown). Despite these precautions, the effective NO-concentrations decreased as a function of the time after addition (inset Fig. 2B). Interaction of NO with the S_1_ state is rather slow and the percentage of centers that is reduced to S_0_ in 1 minute is negligible (G. Schansker, unpublished observation). Unless stated otherwise the samples were bubbled with a NO/N_2_ mixture where the two gasses were mixed in a ratio of 6:14. This results in a concentration in the sample of about 0.6 mM NO, based on the solubility of pure NO of ca 2 mM at 20 °C. However, the effective NO-concentration after 1 min of incubation with NO may have been less (see inset Fig. 2B). In control experiments the samples were bubbled twice with 20 ml N_2_-gas.

**Fig. 2.**
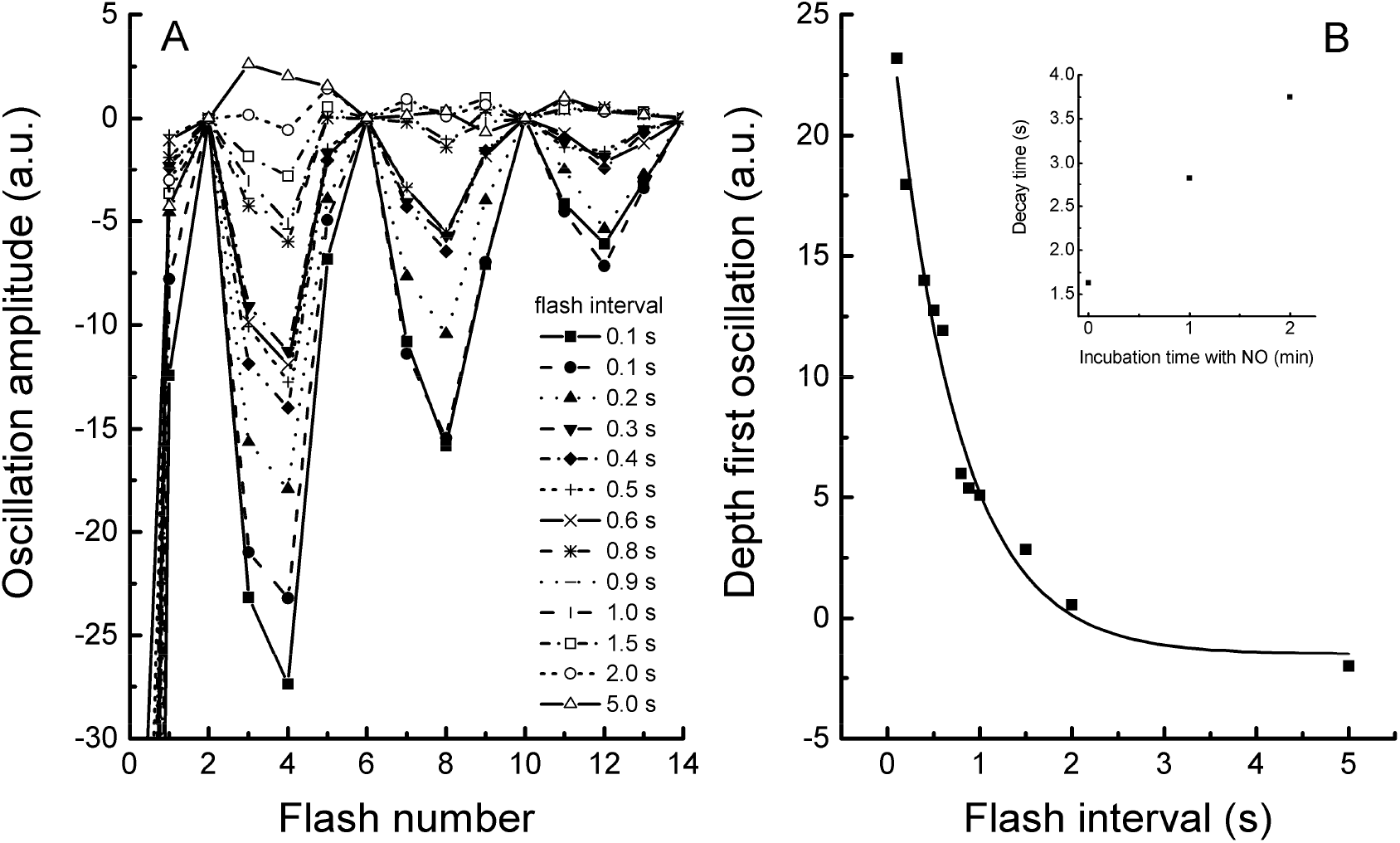
Frequency dependence of the period-4-oscillations in the presence of 0.6 mM NO. A. The measured oscillations and B. The loss of the period-4-oscillations (calculated as the depth of the first oscillation) as a function of the flash frequency. The samples were incubated for 1 minute with NO before the flash series was applied. In the inset of panel B the effect of the incubation time with NO on the decay time (τ) of the S_3_-state is shown.

NO can bind to the non-heme iron of PS II (Diner and Petrouleas 1990) and this may affect the period-4-oscillations by causing acceptor side limitations. Under the experimental conditions, no effect of NO on the electron transfer rates from Q_A_ to Q_B_ (derived from the fluorescence decay kinetics after the xenon-flashes) was observed.

*Hydroxylamine* 50 μM hydroxylamine was used in this study. It was observed that at this concentration hydroxylamine reduced the S_1_ state to S_0_ with a halftime of 1.1 minute (Hanssum and Renger 1985). To minimize this reaction during the experiments the measurement was started immediately after the addition and mixing of the hydroxylamine (in practice about 20 s after addition). Variations in this time may have been a source of some noise in the data.

If hydroxylamine interacts with FeCy in a sample was tested. Addition of hydroxylamine to a cuvette containing 100 μM FeCy had little effect on the size of the FeCy-absorption peak on a minutes time scale, indicating that under my conditions little direct interaction between FeCy and hydroxylamine occurred (data not shown).

### Tyr_D_

In thylakoids, Tyr_D_ is able to donate an electron to the S_2_ and S_3_ states on a seconds time-scale causing a rapid decay of the S_2_ or S_3_ state. In dark-adapted thylakoids this occurs in about 20% of the PS II reaction centers (Vermaas et al. 1984). Both EPR and flash oxygen measurements indicate that in the PS II-membranes used, only a small fraction of Tyr_D_ is in the reduced form. In addition, Tyr_D_ reacts much slower with the S_2_ and S_3_ states in PS II membranes than in thylakoids (Styring and Rutherford 1987). This means that under my experimental conditions the effect of Tyr_D_ on the decay kinetics of the S_2_ and S_3_ states is rather small. No additional pre-flash was given to pre-oxidize Tyr_D_.

## Results

### Frequency dependent loss of period-4 oscillations

The research reported here was initiated by the observation that NO has the ability to eliminate the period-4-oscillations in the F_0_’-level at flash rates of one per second or less. This was indicative for a fast interaction of NO with one or more of the S-states (Schansker and Petrouleas 1998). In Fig. 2A the period-4-oscillations in the F_0_’-level measured at different flash frequencies in the presence of NO are given. The same NO-concentration was used as in Schansker and Petrouleas (1998) but the oscillations were lost with a somewhat slower halftime of 0.5 s (Fig. 2B).

The reason for the differences in the decay times between the data presented here and the data of Schansker and Petrouleas (1998) became clear from the measurements shown in the inset of Fig. 2B. The data show that the decay time (τ) of the S_3_ state increases as a function of the incubation time with NO. From EPR-experiments it is known that the effective NO-concentration declines as a function of the incubation time (the amplitude of the free NO-peak as measured by EPR-spectroscopy declines) (e.g. Goussias et al. 2002). The loss of free NO has been suggested to be due to either dimerization or non-specific reactions (Goussias et al. 2002). Despite the short incubation times used in this study the data in the inset of Fig 2B suggest that a decline in the effective NO-concentration also played a role in the experiments presented here. In earlier studies (Schansker and Petrouleas 1998, Ioannidis et al. 2000) the measurements were made immediately after the addition of NO. Here, the incubation time with NO was increased to 1 min to allow a better equilibration between sample and NO and so to eliminate potential diffusion effects, especially at low NO-concentrations.

The data shown in Fig. 2 have also a practical implication. To study the effects of NO on the various S-states by analyzing the period-4-oscillations high flash frequencies are needed.

### NO-concentration dependence

In a previous paper (Ioannidis et al. 2000) the S-states that could interact rapidly with NO were identified. There, it was observed that the decay times for the S_2_ and S_3_ decay could be accelerated considerably in the presence of 0.6 mM NO: from 150 s to 2.1 s for the S_2_ to S_1_ decay and from 28 s to 0.4 s for the S_3_ to S_1_ decay. In Fig. 3 the NO-concentration dependencies of the reaction rates of the S_3_ (main figure) and S_2_ (inset) states are shown. The S_3_-decay rate increases considerably as a function of the NO-concentration. The reaction rate increased in a sigmoidal way in response to the NO-concentration. This could indicate that at least 2 NO molecules, acting in a cooperative or sequential fashion, are needed for the formation of the S_1_-state out of the S_3_-state. In the case of the S_2_-state a similar increase of the decay rate as a function of the NO-concentration was observed. The incubation time with NO has also an important effect on the reaction rates. In the case of the S_3_-state the sigmoidal NO-concentration dependence is shifted to lower NO-concentrations by reducing the incubation time with NO. For the S_2_-state the incubation-time effects are more complex. For samples measured after 1 min incubation with NO, the NO-concentration dependence has a sigmoidal character, whereas for samples measured immediately after the addition of NO the response to changes in the NO-concentration is more exponential than sigmoidal. At the same time, it is worth noting that at the lowest NO-concentration, 1 min incubation with NO leads to a faster reaction rate indicating that under these conditions there is an interplay between diffusion limitations and the incubation-dependent decline of the effective NO-concentration. For the measurement of the decay rates of the S_2_ and S_3_ states immediately after the addition of NO, a smaller data set was available. Even so, the trends can be quite clearly observed.

**Fig. 3.**
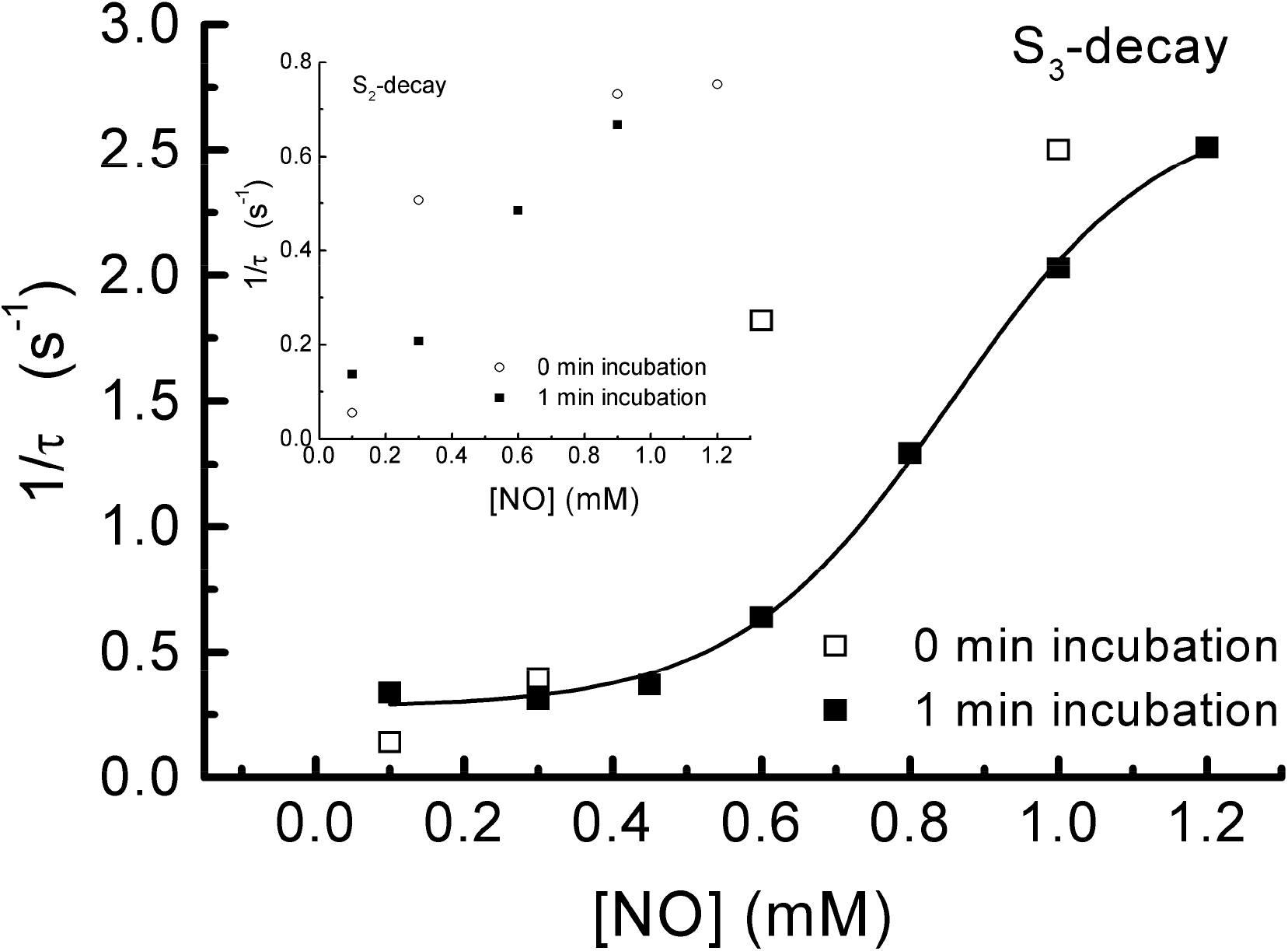
NO-concentration dependence of the S_3_ decay rates. The effect of the incubation time with NO on the S_3_ decay rate is also shown. In the inset a similar data set for S_2_ decay rates is shown. Each decay rate is based on 12-15 measurements. The decay rate for the lowest NO-concentration represents the decay to the S_2_-state. Standard deviations are similar to those shown in the inset of Fig. 5 at the appropriate pH.

In Fig. 4 the fluorescence spectra of the S_3_-decay-kinetics obtained with 0.1, 0.8 and 1.2 mM NO are compared. At 0.1 mM of NO the S_3_-state decays via the S_2_-state to S_1_ whereas at 0.8 and 1.2 mM no S_2_-intermediate can be discerned (see Ioannidis et al. (2000) for the interpretation of the spectra). It was observed that at 0.3 mM NO still 10-15% S_2_-intermediate can be discerned but at concentrations above 0.3 mM NO this component totally disappears and the S_3_ decay becomes for all practical purposes monophasic. Fitting the 0.1 mM NO data using sequential kinetics indicated that at intermediate times about 50% S_2_ accumulates. If Figs. 4B (0.8 mM NO) and 4C (1.2 mM NO) are compared, it can be observed that at intermediate times (almost equal amounts of S_3_ and S_1_) at a concentration of 0.8 mM NO the trace is almost flat with a slight deviation on the right-hand side. This deviation is stronger in the presence of 1.2 mM NO. Under those conditions, a trace that looks like an S_0_-trace becomes visible (compare with Fig. 1D). This could indicate that at high NO-concentrations some S0 may be formed, be it with a low probability.

**Fig. 4.**
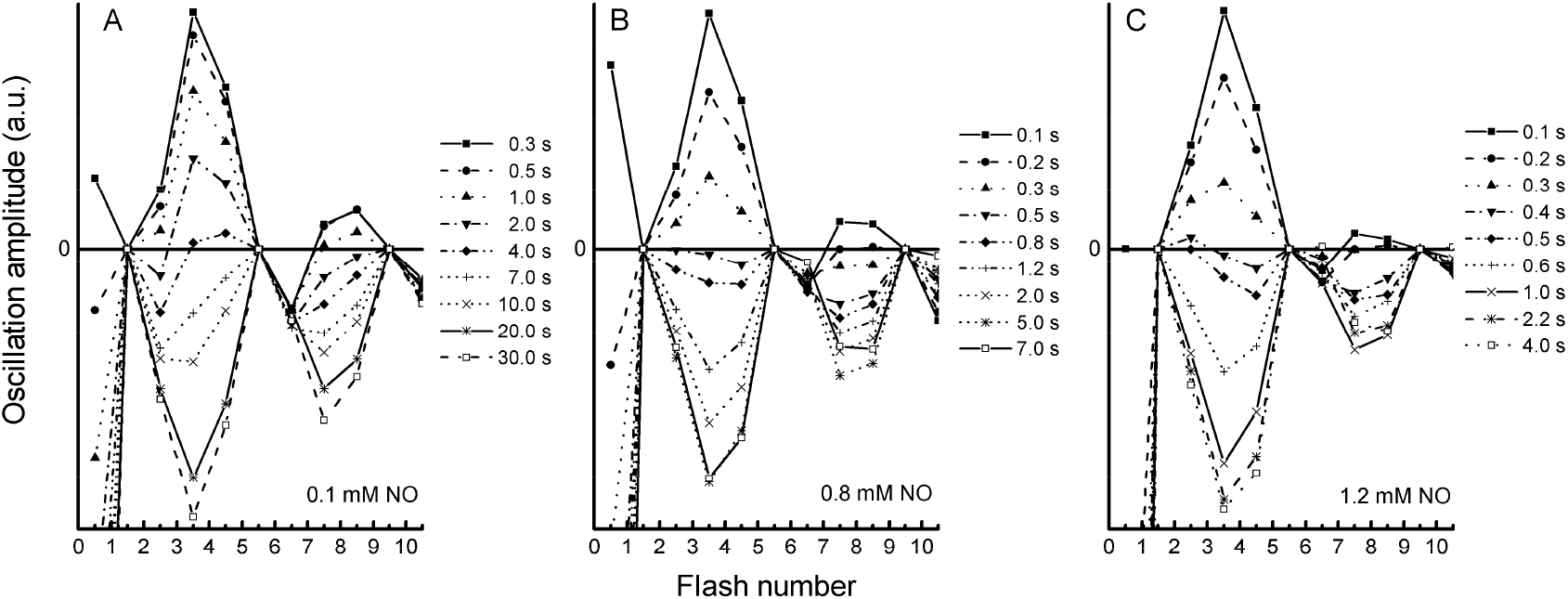
Period-4-oscillations as a function of the dark-interval time between the pre-flashes and the flash series. The samples were incubated for 1 min with 0.1 mM NO (A), 0.8 mM NO (B) and 1.2 mM NO (C) before the start of the measurements. Representative data sets are shown.

The fact that it is still possible to assay the period-4-oscillations after the NO-treatment indicates that PS II can still turnover normally after the S_2_ and S_3_ states have interacted with NO. To demonstrate this, the decay time of the S_2_ state in the presence of NO starting from a dark-adapted sample or from a sample to which two pre-flashes had been given was determined. The flash to produce the S_2_-state was given 2 s (1.2 mM NO) or 4 s (0.6 mM NO) after the two initial flashes were given. In both cases the decay time of the S_2_ state was unaffected by an earlier reaction of the S_3_-state with NO (data not shown). This is an important observation because it shows that the reduction of the S_3_-state by NO does not affect the functional integrity of the oxygen-evolving complex.

### pH-dependence of the S_3_ – NO interaction

The pH-dependence of the S_3_-decay may make the involvement of a protonation in the decay process visible. In Fig. 5 the pH-dependence of the interaction between NO and the S_3_-state is shown. It is observed that the reaction rate of NO with the S_3_ state between pH 6.0 and 7.7 decreases sigmoidally with a pK-value of 6.9. Above pH 7.7 the data seem to indicate that the rate starts to decline again. However, this is only one point in a pH-range where the donor side of PS II in PS II-membranes starts to become unstable. On the other hand, the result was quite reproducible. Between pH 6 and 7.7 the reaction rate decreased by a factor 2.2. In Table 1 the S_3_-decay rates at pH 6.5 and pH 7.5 in the presence of 0.6 mM NO are compared with those in the presence of 1.2 mM. At the higher NO concentration the ratio rate(pH 6.5)/rate(pH 7.5) increases to 3.8. This indicates that a full protonation may occur on going from S_3_ to S_1_, but this is masked to some extent by the fact that the protonation step is only partially rate limiting.

**Fig. 5.**
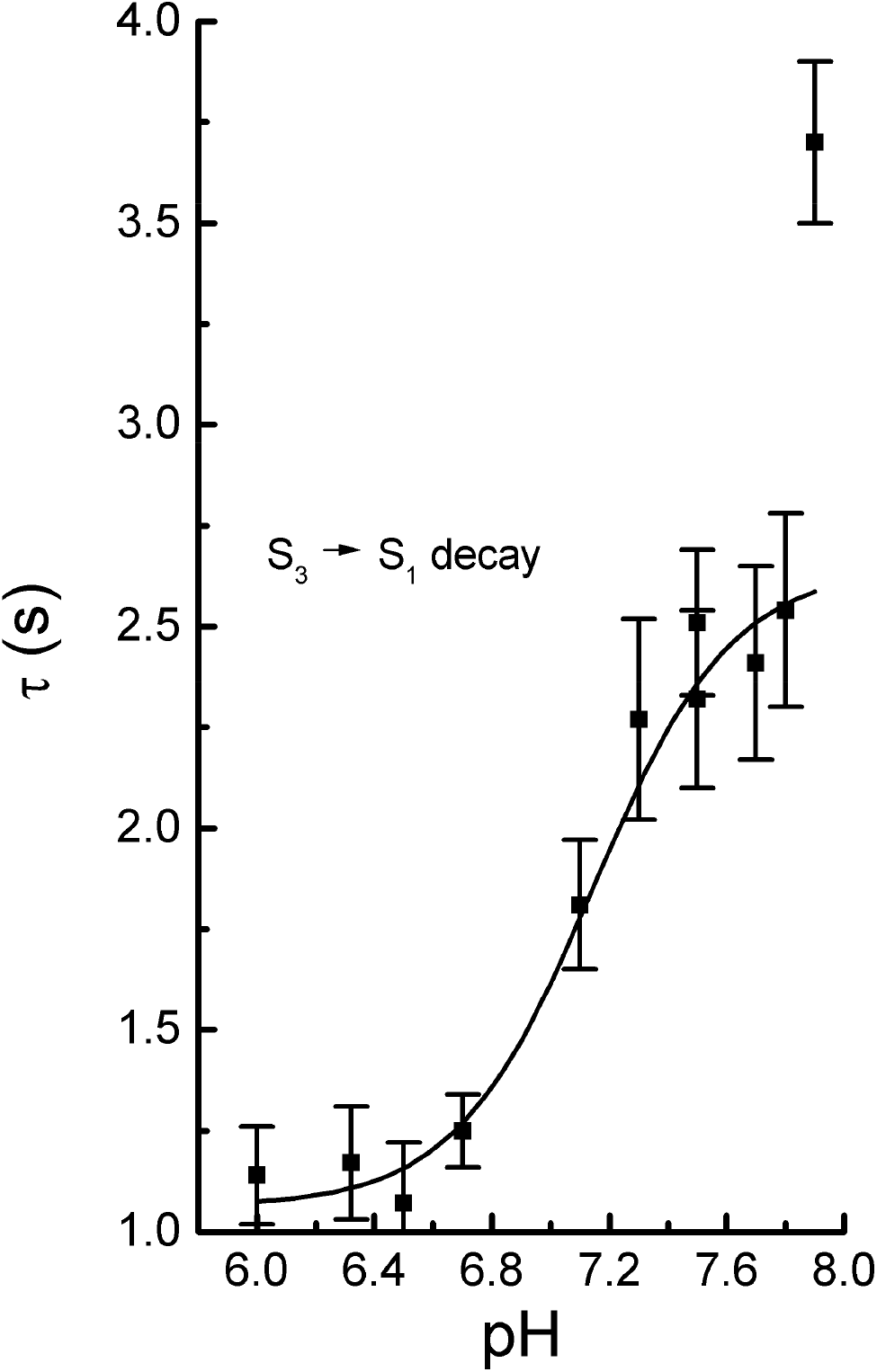
pH-dependence of the S_3_-decay rate in the presence of 0.6 mM NO. The samples were incubated for 1 min with NO before the start of the measurement. In the inset the S_3_-decay times (τ) and their standard deviation have been plotted. Each decay rate is based on 12-15 measurements.

**Table 1.** Effect of the NO-concentration and pH on the decay rates of the S_3_-state. Each decay rate is based on 12 to 15 measurements.

|  | NO-concentration |  |
| --- | --- | --- |
|  | 0.6 mM NO | 1.2 mM NO |
| Decay rate ($s^{-1}$) (pH 6.5) | 0.9 | 2.5 |
| Decay rate ($s^{-1}$) (pH 7.5) | 0.5 | 0.7 |
| ratio(pH6.5/pH7.5) | 2.0 | 3.8 |

Fig. 5 demonstrates that at pH-values between 6 and 7 the standard deviation is quite small. At higher pH-values the standard deviation increases because the amplitude of the period-4 oscillations declines.

### Comparison between NO and hydroxylamine reactivity

The observations on the decay-time of the S_3_ state seemed to contradict measurements of Messinger and Renger (1990) using hydroxylamine and hydrazine. Messinger and Renger (1990) demonstrated that hydrazine is able to accelerate the decay of the S_2_-state considerably whereas the decay of the S_3_ state in the presence of hydrazine is hardly faster than in control samples. Although the authors did not show the data, they wrote that the effect of hydroxylamine was comparable to that of hydrazine. The experiments presented by Messinger and Renger (1990) were repeated using the one-electron donor hydroxylamine. In Fig. 6 a comparison is made between the decay kinetics of the S_2_ and S_3_ states in the presence of either hydroxylamine or NO. As Fig. 6 demonstrates the results presented here confirmed the measurements presented by Messinger and Renger (1990). In the presence of hydroxylamine, the decay of the S_2_ state is accelerated considerably. Decay times are comparable to those in the presence of NO. On the other hand, the decay-time of the S_3_ state in the presence of hydroxylamine was closer to the decay time in control samples. The decay time of the S_3_ state in the presence of hydroxylamine determined from the data in Fig. 6 may underestimate the true value somewhat. Both the S_2_ to S_1_ and the S_1_ to S_0_ transition in the presence of hydroxylamine were faster than the S_3_ to S_2_ decay. As a consequence, the last part of the S_3_ decay is lost in the intermediate mixture of states that is formed. On a time-scale of several minutes the S_-1_ state started to build up as a more or less final reduction state. It has to be noted that the hydroxylamine concentration of 50 μM is very low compared to the estimated NO-concentration of 1.2 mM. Since the decay times of the S_2_-state are the same this indicates that hydroxylamine is a more efficient reductant of the Mn-cluster than NO. This is strengthened by the observation that hydroxylamine is a quite efficient reductant of the S_1_-state. Hanssum and Renger (1985) observed an almost full reduction from S_1_ to S_0_ within 5 minutes after the addition of NH_2_OH and Messinger et al. (1991) observed a 50% reduction of S_1_ during the same period. In both cases 50 μM NH_2_OH was used. NO, in contrast, seems to have little effect on the S_1_ state on a minutes time-scale (G. Schansker, unpublished observation).

**Fig. 6.**
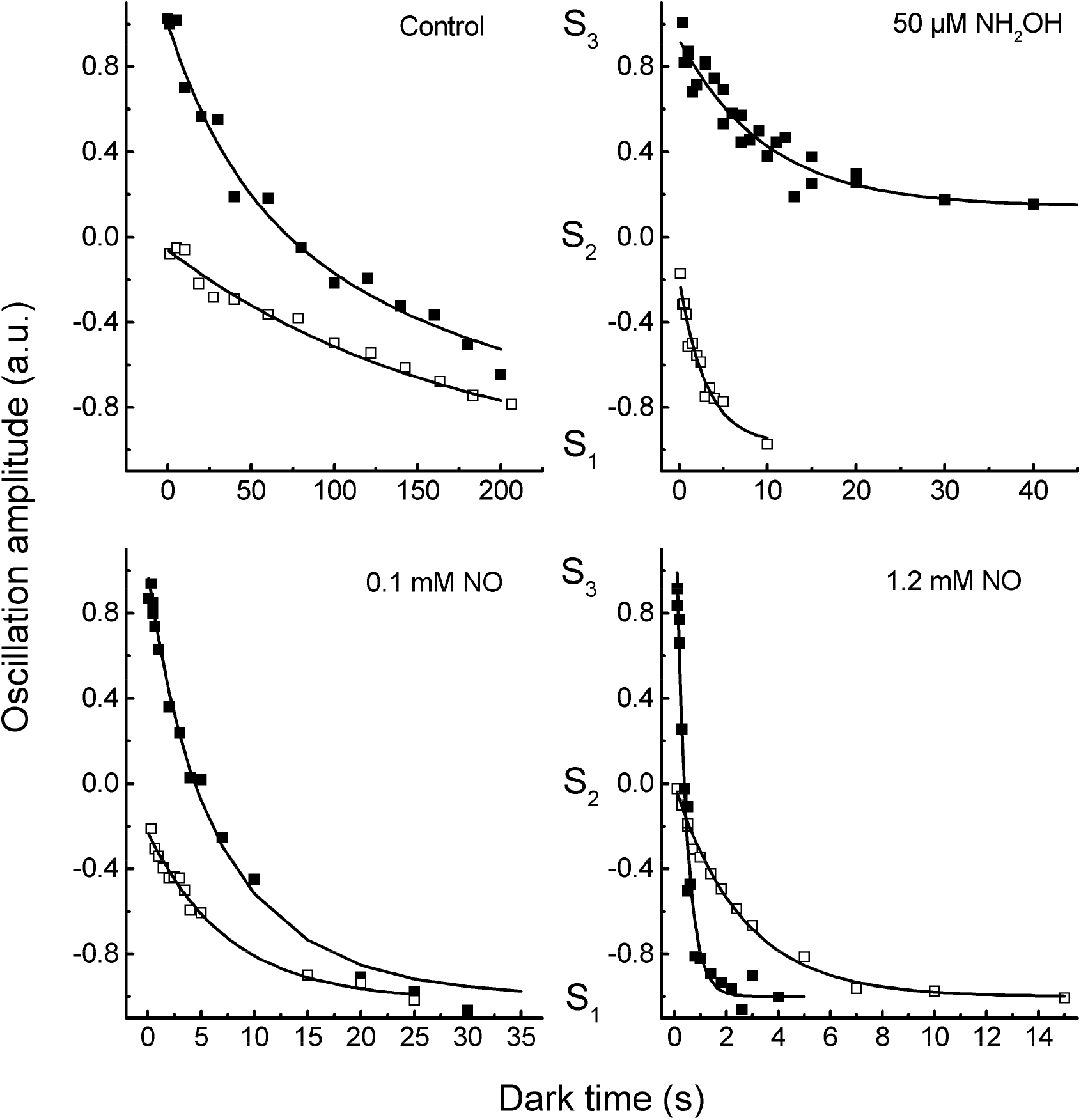
Comparison of the decay kinetics of the S_2_ and S_3_ states. A. In the absence of electron donors and in the presence of 50 μM hydroxylamine (B), 0.1 mM NO (C) and 1.2 mM NO (D). In the panels the S_3_ state was set to 1, the S_2_ state was set to 0 and the S_1_ state was set to −1. Representative data sets are shown.

With respect to the S_2_-decay in the presence of hydroxylamine it has to be noted that the changes in the period-4 oscillations for this decay process were rather limited. However, the correctness of the decay data is supported by the fact that the determined decay rate is very similar to that determined earlier using another technique (Messinger and Renger 1990). Further support comes from the fact that a good dataset is obtained (see the NH_2_OH-panel of Fig. 6)

## Discussion

### Flash fluorescence technique

The first observations of period-4 oscillations in the fluorescence signal were made almost 50 years ago (Joliot et al. 1971, Delosme 1971, Zankel 1973). If the fluorescence technique is compared with the use of the bare oxygen electrode the last technique has one important advantage: the interpretation of the measurements is straightforward. Oxygen is only produced on going from S_3_ via S_4_ to S_0_. The data from the fluorescence approach are more difficult to interpret, but lower chlorophyll concentrations can be used, the response time is much shorter (it is flash frequency-dependent which means in this case a time resolution of 100 ms) and there are no problems due to interactions of electron acceptors or donors with the oxygen electrode.

The origin of the period-4-oscillations in the F_0_’-level has not yet been resolved. In an earlier paper (Schansker et al. 2002) some literature observations were quoted on S-state-dependent proton release and the protonation-state of nearby antenna-molecules. Jahns and Junge (1992) observed that part of the antenna buffers proton release by the oxygen-evolving complex in a DCCD-dependent way and Delrieu (1998) observed that the presence of a full complement of LHCII-antenna proteins was necessary for strong oscillations. In this study it was observed that there was a clear relationship between the quality of the PS II-membrane preparations and the depth of the period-4-oscillations (cf. Schansker and Petrouleas 1998 with Fig. 2A).

### Concentration-dependence

In Figs. 3 and 4, three observations attract attention. In the first place, it is observed that for NO-concentrations >0.3 mM the S_3_ to S_1_ decay in the presence of NO was faster than the S_2_ to S_1_ decay (see also Ioannidis et al. 2000). In the second place, it is observed that at NO-concentrations <0.3 mM an S_2_-intermediate is observed (at 0.1 mM 50% S_2_ is build up at intermediate times). And in the third place, the [NO]-dependence of the S_3_-decay is sigmoidal. These observations suggest either that 2 NO-molecules interact in a concerted fashion with the S_3_-state or, and this seems more likely, that the second NO-molecule reacts with a highly reactive intermediate created by the reaction of the first NO-molecule with the S_3_-state. In the absence of a second NO-molecule the S_2_-state is formed. It may be noted that if the interaction of NO with the S_3_ state would lead to a modified S_2_ state (S_2_*) (see e.g. Ioannidis and Petrouleas 2002) it has to be assumed that S_2_* cannot be distinguished from S_2_ on the basis of the assay. And secondly, that if the second NO-molecule reacts with an intermediate state the concentration of this intermediate state remains below the detection limit of the assay.

### pH-dependence

In Fig. 5 the pH-dependence of the decay rates of the S_3_ state in the presence of 0.6 mM NO is shown. In the range studied, one sigmoid with a pK-value of 6.9 is observed. Between pH 6.0 and 7.7 the decay rate of the S_3_-state decreases by a factor of 2.2 (see Table 1). This effect is stronger at higher NO-concentrations (3.8 at 1.2 mM NO). Since the reaction rates are faster at 1.2 mM, the increase to 3.8 seems to indicate that a protonation is probably involved in the S_3_-decay but that it is only partially rate limiting.

The data in Fig. 5 allow a comparison with those presented by Messinger and Renger (1994). These authors used the interaction of the S_3_-state with two internal reactants, Q_B_^−^ and Tyr_D_ to determine the pH dependence of the S_3_ state. In both cases the internal reactants do not interact directly with the S_3_-state. The pH-dependences of these two reactions differed in several respects. The recombination of Q_B_^−^ with the S_3_-state slowed down over the whole pH-range measured (4.5-8.0) whereas the interaction with Tyr_D_ was relatively slow around pH 6.5 and accelerated at both higher and lower pH-values. A comparison of these data with the pH-profile of the interaction of the S_3_-state with NO is useful because NO interacts directly with the S_3_-state and there is little indication of an effect of pH on NO-chemistry in the pH-range measured (Gross and Wolin 1995). The comparison indicates that especially for the interaction of Tyr_D_ with the S_3_-state the observed pH-profile is a superimposition of pH-effects on the S_3_-state, on the electron transfer route followed and on Tyr_D_. It has to be noted that the reaction time of the S_3_-state with NO was faster than the reaction time with Tyr_D_ over the whole pH-range measured. On the other hand, the pH-profile of the interaction of the S_3_-state with NO may be comparable to that of the recombination reaction between Q_B_^−^ and the S_3_-state (Messinger and Renger 1994) but the time resolution of the published data is not high enough to determine this with certainty. Messinger and Renger (1994) argued that the pH-profile of the interaction of Tyr_D_ with the S_2_ and S_3_ states would be mainly determined by the properties of these S states. The data presented here indicate that this may not be valid and that in this respect Tyr_D_ is not a good probe for the properties of the S states.

### NO, iodide and hydroxylamine

Comparing the reactivity of NO towards the S_3_-state with that of other compounds there is one important difference. Only the interaction with NO leads to a near monophasic decay to the S_1_-state. Hydroxylamine is not reactive towards the S_3_ state (Messinger and Renger 1990) and Wincencjusz et al. (1999) suggested that iodide is also able to donate an electron to the S_3_-state but in that case the reaction resulted in the S_2_-state.

### Proposed mechanism for the interaction of NO with the S_3_-state

Analyzing the data, the following global reaction mechanism can be proposed: NO donates an electron to the S_3_-state, this induces a reactive intermediate state and then two things can happen. At high NO concentrations a second NO molecule can react with this intermediate state leading to a fast decay to the S_1_ state (Figs. 2, 4 and 6). At low NO concentrations the intermediate state decays to the S_2_ state and then relatively slowly to S_1_ (Figs. 4 and 6). The pH dependence of the interaction of NO with the S_3_ state indicates that the reaction involves a protonation (Table 1 and Fig. 5). At the same time the data confirm that hydroxylamine did not really accelerate the S_3_ decay rate (Fig. 6) as observed before by Messinger and Renger (1990). It is also important to note here that the assay used is a functional assay, monitoring the fluorescence quenching effect of the different S-states on the F_0_ level. In other words, the states induced by NO behave as normal S-states. There is no indication of damage to the Mn-cluster due to NO.

Twenty years ago, the possibility that the S_3_ state had a radical character was still considered a viable possibility (Styring and Rutherford 1988, Lavergne 1991, Roelofs et al. 1996, Ioannidis et al. 2000, Messinger et al. 2001, Yano and Yachandra 2007) and it is known that NO interacts better with radicals (e.g. Eiserich et al. 1995, Ford 2004) than with metals (e.g. Ford 2004). The difference in the decay rate of the S_2_ state in the presence of, respectively, hydroxylamine and NO (Fig. 6), which is an reduction of Mn(IV) to Mn(III) (e.g. reviewed by Dau and Haumann 2008, Sauer et al. 2008), confirms that hydroxylamine is the more effective reagent in this respect. However, the consensus is at the moment that on each S-state transition a Mn-molecule becomes oxidized (Dau et al. 2012, Siegbahn 2012, Yano and Yachandra 2014, Vinyard and Brudvig 2017, Junge 2019, Lubitz et al. 2019). In the second place, hydroxylamine is thought to be a structural mimic of water (Hanssum and Renger 1985). Taking all these things together and considering as well our much better understanding of the S-states and S-state transitions, is it then possible to come up with a more mechanistic description of the interaction of NO with the S_3_ state? Based on density functional theory (DFT) calculations making use of the 1.9 Å PS II structure of Shen and coworkers (Umena et al. 2011) the following reaction sequence for the S_2_ to S_3_ transition was obtained: following a charge separation Tyr_Z_ is oxidized by P680^+^; the positive charge on TyrZ leads to the release of a proton into the lumen. Tyr_Z_^+^ then oxidizes a Mn(III) to Mn(IV) – the Mn that is designated in the literature as Mn1 – which is accompanied by a tighter binding of a water molecule to Mn1 and a deprotonation of this water molecule, followed by several proton transfers (Siegbahn 2012). It could then be imagined that NO can partially reverse this process, where the tightly bound water molecule would get an electron from NO (and a proton, Fig. 5) leading to an intermediate state sensitive to a second NO molecule. Hydroxylamine, as a structural water mimic, cannot do this and, as the experiments show, is ineffective when it comes to interacting with the S_3_ state.

## Acknowledgements

GS is indebted to dr. Vasili Petrouleas for useful discussions as well as practical suggestions and for offering him the opportunity to carry out the present study and to dr. Nikos Ioannidis for a critical reading of an earlier version of the manuscript. He would like to thank dr. DM Kramer for the instructions needed to construct the fluorescence apparatus, mr. Panos Vassilopoulos for the construction of the fluorescence apparatus and mr. Laszlo Sasz for writing the final version of the operating software. Financial support by the EC TMR program ERBFMRXCT980174 is gratefully acknowledged.

## Abbreviations

Chl: chlorophyll
EXAFS: extended X-ray absorption fine structure
FeCy: ferricyanide, K_3_[Fe(CN)_6_]
F_0_: fluorescence intensity when all PS Il-reaction centers are open, F_0_’, initial fluorescence level
HEPES: N-(2-hydroxyethyl) piperazine-N’-2-ethane-sulphonic acid
LED: light emitting diode
MES: 2-(*N*-morpholine)-ethanesulfonic acid
NO: nitric oxide
PS II: photosystem II
PS II membranes: thylakoid membrane fragments enriched in PS II
Q_A_ and Q_B_: the primary and secondary plastoquinone electron acceptors of PS II, S states (S_0_, …, S_4_), oxidation states of the oxygen-evolving complex
Tyr_D_: slow tyrosine electron donor of PS II

